# Transforming Insect Population Control with Precision Guided Sterile Males

**DOI:** 10.1101/377721

**Authors:** Nikolay P. Kandul, Junru Liu, Hector M. Sanchez C., Sean L. Wu, John M. Marshall, Omar S. Akbariut

## Abstract

The sterile insect technique (SIT) is an environmentally safe and proven technology to suppress wild populations. To further advance its utility, a novel CRISPR-based technology termed “precision guided SIT” (pgSIT) is described. PgSIT mechanistically relies on a dominant genetic technology that enables simultaneous sexing and sterilization, facilitating the release of eggs into the environment ensuring only sterile adult males emerge. Importantly, for field applications, the release of eggs will eliminate burdens of manually sexing and sterilizing males, thereby reducing overall effort and increasing scalability. To demonstrate efficacy, we systematically engineer multiple pgSIT systems in *Drosophila* which consistently give rise to 100% sterile males. Importantly, we demonstrate that pgSIT-generated males are fit and competitive. Using mathematical models, we predict pgSIT will induce substantially greater population suppression than can be achieved by currently-available self-limiting suppression technologies. Taken together, pgSIT offers to transform our ability to control insect agricultural pests and disease vectors.

## Introduction

CRISPR-based genome editing has revolutionized the capacity for precise genome manipulations in nearly every organism studied (reviewed in ^1^). For example, recently it has been used to develop extremely efficient homing based gene drives that can bias Mendelian inheritance rates with up to 99% efficiency in many animals including flies, mosquitoes, and mice ^2–5^, revolutionizing an entire new field termed “Active Genetics ^6^.” While these innovative technologies bear the potential to provide worldwide solutions to combat vector-borne diseases, improve agriculture and control invasive species, ongoing discussions are underway to define mechanisms of governance to ensure the technology is ethically, and safely, developed and implemented ^7–9^. Notwithstanding, current drive designs are limited by the rapid evolution of resistance ^10^, and therefore future research is necessary to develop drives that can limit and overcome evolved resistance ^11,12^. While these discussions and developments are advancing, given the precision, simplicity, and efficiency of CRISPR, we aimed to develop a novel, safe, controllable, non-invasive, genetic CRISPR-based technology that could be transferred across species and implemented worldwide in the short-term to combat wild populations.

Coined independently by Serebrovskii, Vanderplank, and Knipling, mass-production and release of sterile males, known as the Sterile Insect Technique (SIT), has historically been used to control, and eradicate, insect pest populations dating back to the mid-1930s ^13–17^. Traditional methodologies have relied on DNA-damaging agents for sterilization, substantially reducing overall fitness and mating competitiveness of released males. To overcome these issues, microbe-mediated infertility techniques such as *Wolbachia*-based incompatible insect technique (IIT) *^18,19^*, and modern genetic SIT-like systems such as the Release of Insects carrying a Dominant Lethal (RIDL)^20^, and other methodologies to release fertile males that genetically kill females such as female-specific RIDL (fsRIDL)^21^, and autosomal-linked X-chromosome shredders ^22^ have been developed (reviewed in ^23^). While these first-generation genetic SIT technologies represent significant advances, IIT strictly requires no infected females to be released which is difficult to achieve in the field, and the use of tetracycline known to ablate the microbiota ^24^ compromises the fitness of RIDL/fsRIDL males, and X-chromosome shredders can in principle only be developed in species with heterogametic sex chromosomes, thereby limiting wide applicability to other species. Therefore, it would be logistically advantageous to employ more efficient SIT-based technologies that could be deployed as eggs by which only sterile males would survive, to date such optimal genetic technologies do not exist.

Here we develop a next-generation highly-efficient precision guided SIT (pgSIT) technology that can be deployed as eggs which only give rise to sterile males. PgSIT functions by exploiting the precision and accuracy of CRISPR to simultaneously disrupt genes essential for either female viability or male fertility. It utilizes a simple breeding scheme requiring two homozygous strains -one expressing Cas9 and the other expressing double guide RNAs (*dgRNAs*). A single mating between these strains mechanistically results in synchronous RNA-guided dominant biallelic knockouts of both target genes throughout development, resulting in the complete penetrance of desired phenotypes in all progeny (Fig. 1A). We show that pgSIT is extremely robust at genetically sexing and simultaneously sterilizing resulting progeny reproducibly with 100% efficiency. Moreover, we demonstrate that pgSIT sterile males are fit and can compete for mates. Taken together, pgSIT offers to lead far superior population suppression over existing approaches, thereby revolutionizing SIT-mediated control of wild populations.

**Fig. 1.**
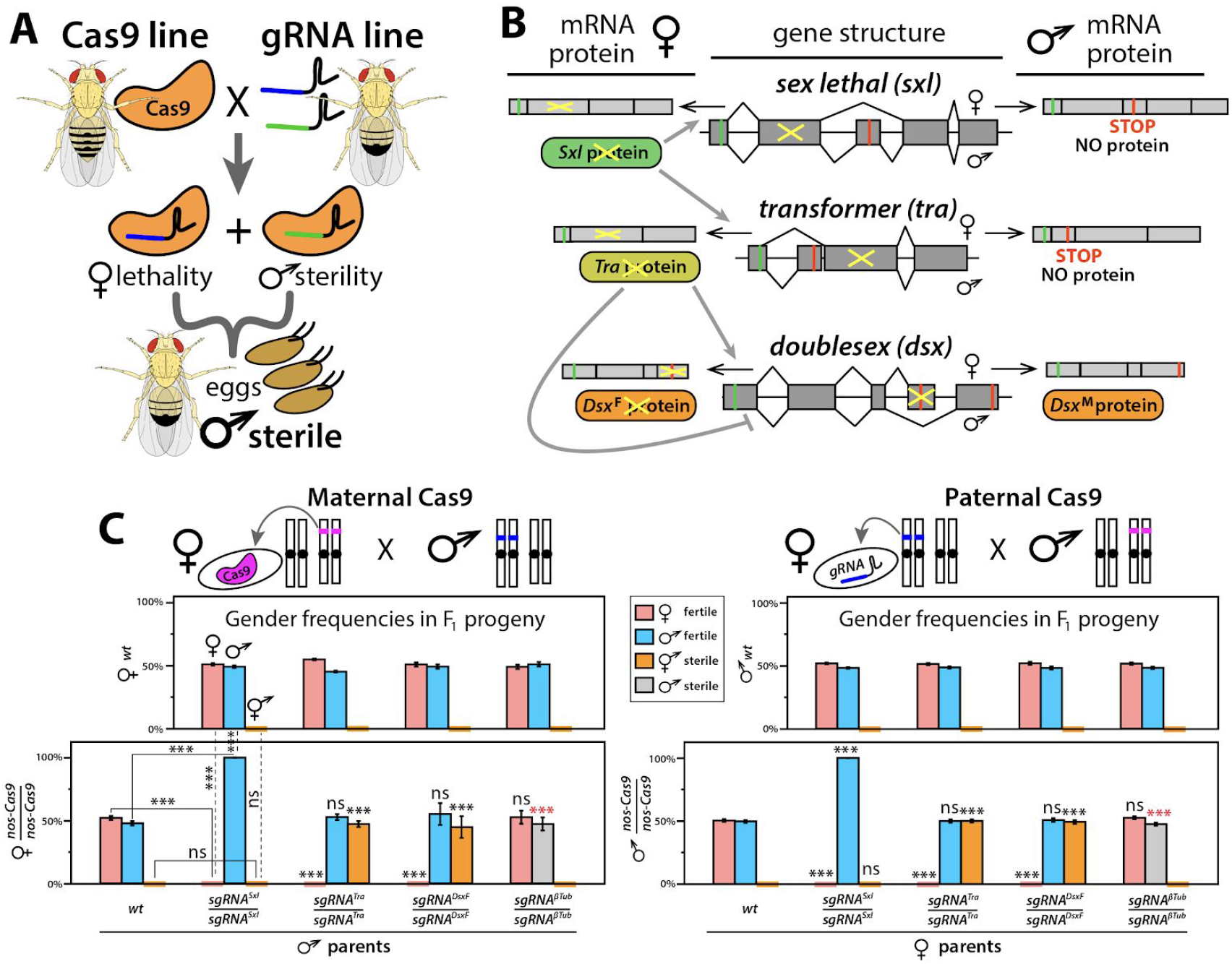
Schematic representation of precision guided Sterile Insect Technique (pgSIT), including gene targets and assessment of single guide RNAs (sgRNAs). (**A**) A schematic of pgSIT utilizing two components of the binary CRISPR/Cas9 system - Cas9 and gRNAs-maintained as separated homozygous lines, their cross results in simultaneous knockouts of two genes required for female viability and male sterility resulting in survival of only F_1_ sterile males. (**B**) A schematic of sex specific alternative splicing in *sxl, tra* and *dsx* regulated by female expression of Sxl and Tra proteins (gray lines) (modified from ^59^). Disruption of female-specific exons of key sex-determination genes, *sxl, tra* and *dsx*, disrupts female development. PgSIT exon targets indicated by yellow crosses. (**C**) Bar graphs of average gender frequencies in F_1_ progeny. Two top panels depict gender frequencies from bidirectional control crosses of homozygous sgRNA lines to wild type (*wt*) indicating that both fertile females and males (♀ and ♂) are present at similar ratios, but no sterile intersexes (⚥) were identified. Bottom two panels show gender frequencies from crosses of homozygous *nanos-Cas9* (*nos-Cas9*) to *wt* (control) and four homozygous sgRNA lines (experiment). Independent of maternal or paternal Cas9 inheritance, 100% of trans-heterozygous *sgRNA^Sxl^* ♀ died, 100% of trans-heterozygous *sgRNA^Tra^* and *sgRNA^DsxF^* ♀ were masculinized into sterile intersexes ⚥, and 100% of trans-heterozygous *sgRNA^βTu^* ♂ were sterile. Gender frequencies and fertility in trans-heterozygotes were compared to those in corresponding progeny of control crosses with *Cas9* (solid lines) or *sgRNAs* (dashed lines) and *wt* flies. Each bar shows an average gender frequency and one standard deviation. Statistical significance was calculated with a *t* tests assuming unequal variance, and for male sterilization, *P* values were calculated using Pearson’s Chi-squared test for contingency tables (red*). (*P* > 0.001***).

## Results

### Binary CRISPR Induced Female Masculinization/Lethality, or Male Infertility

To engineer pgSIT, we first generated single guide RNA (sgRNA) and spCas9 (Cas9 from hereon) expressing lines in *Drosophila*. In total nine homozygous sgRNAs lines were developed to target genes essential for female viability, or genes important for male fertility. For female viability, these genes included sex-specifically alternatively spliced sex-determination genes including sex *lethal (Sxl*, two separate transgenic lines - *sgRNA^Sxl^, sgRNA^Sxl-B^), transformer (tra*, two separate lines - *sgRNA^Tra^, sgRNA^Tra-B^*) or *doublesex* (*dsxF, sgRNA^DsxF^*) (Fig. 1B, S1) ^25–28^. To disrupt male fertility, we targeted genes active during spermatogenesis, such as *βTubulin 85D (βTub, sgRNA^βTub^)^29^, fuzzy onions (fzo, sgRNA^Fzo^)^30^, protamine A (ProtA, sgRNA^ProtA^)^31^*, or *spermatocyte arrest (sa, sgRNA^Sa^)^32^* (Fig. S1). To promote robust Cas9 expression, we established three homozygous Cas9 expressing lines under control of two strong predominantly germline specific promoters, including *nanos* (*nos-Cas9*) or *vasa* (*vas-Cas9*)^33,34^, and one ubiquitous promoter to enable robust expression in both somatic and germline tissues during nearly all developmental life stages, *Ubiquitin 63E (Ubi-Cas9)^35^*(Fig. S2). Downstream (3’) to the promoter-driven Cas9 we included a self-cleaving T2A peptide and eGFP coding sequence, together serving as a visual indicator of promoter activity ^36^ (Fig. S1, S2).

To assess the genetic activity of the sgRNA lines, we crossed each strain to *nos-Cas9*, and examined resulting trans-heterozygous F_1_ progeny. From these crosses, 4/9 of the sgRNAs, including *sgRNA^Sxl^, sgRNA^Tra^, sgRNA^DsxF^, sgRNA^βTub^*, displayed expected phenotypes and were subjected to further characterization. To further evaluate these four sgRNAs, we bidirectionally crossed them to wild type *wt* (# progeny (n) =3519, # replicates (N) = 24), or to homozygous *nos-Cas9* (n=3628, N=28)(Table S1). As expected, the *wt* crosses produced no significant gender ratio deviations or compromised fertility (n=4371, N=30) (Fig. 1C, Table S1). Interestingly however, regardless of whether *nos-Cas9* was maternally or paternally inherited, all F_1_ trans-heterozygotes inheriting *sgRNA^Sxl^* were 100% male (n=540, N=7), and 100% of trans-heterozygous females inheriting *sgRNA^Tra^* or *sgRNA^DsxF^* were converted into sterile masculinized intersexes unable to oviposit eggs (n=942, N=14), and 100% of *sgRNA^βTub^* trans-heterozygous males were sterile (n=517, N=7)(Fig. 1C, Table S1). These phenotypes were moleculary explored at the targeted genetic loci, and as expected we discovered that all sequenced flies (n =16) had mosaic insertions/deletions (indels) precisely at the targeted loci (Table S2).

### Creation of Populations of 100% Sterile Males

The goal of pgSIT is to disrupt genes essential for male fertility/female viability simultaneously to ensure that all surviving F_1_ offspring are sterile males. To achieve this feat, leveraging the results from above, we generated three additional homozygous strains expressing multiplexed double gRNA (*dgRNA*) combinations, including *dgRNA^βTub-Sxl^, dgRNA^βTub-Tra^*, and *dgRNA^βTub-DsxF^* (Fig. S1). To genetically assess the activity of these pgSIT strains, we bidirectionally crossed each line to *w*t, or homozygous Cas9 (either *nos-Cas9, vas-Cas9*, or *Ubi-Cas9*). As expected, the *wt* crosses produced no significant gender deviations or compromised fertility (n=5747, N=36) (Fig. 2A, Table S3). Interestingly however, the crosses between *dgRNA^βTub-Sxl^* with each Cas9 strain resulted in 100% female lethality due to disruption of *sxl*, in addition to 100% male sterility due to simultaneous disruption of *βTub* (n= 2521, N=24) (Table S3). Moreover, 100%) females from crosses between each Cas9 strain and *dgRNA^βTub-Tra^* (n=1697, N=24) or *dgRNA^βTub-DsxF^* (n=1791, N=24) were masculinized into sterile intersexes due to disruption of either *tra* or *dsx*, and 100% male offspring were sterile due to simultaneous disruption of *βTub* (n=4231, N=48). These findings demonstrate that the ability to form highly active Cas9-gRNA complexes was not saturated by dgRNAs and the pgSIT approach works reproducibly with unprecedented efficiency (Fig. 2A-B, Table S3).

**Fig. 2.**
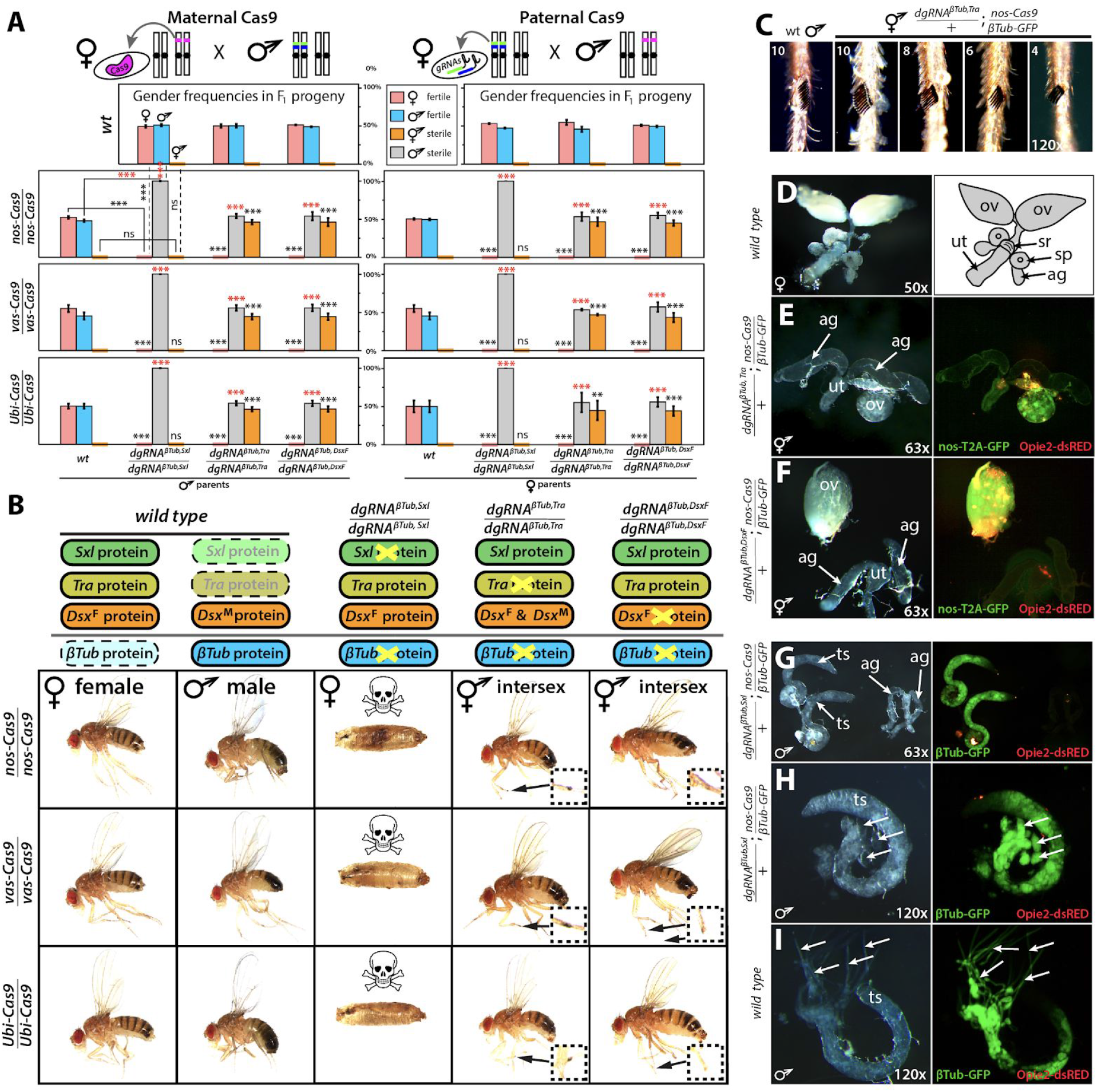
Development and characterization of multiple highly efficient pgSIT systems. (**A**) Gender (♀, ♂, and ⚥) frequencies of trans-heterozygous F_1_ progeny resulting from crosses between double gRNAs (*dsRNA*) and *Cas9* homozygous lines. Three dgRNAs, each targeting *sxl, tra* or *dsx* combined with *βTub*, were bidirectionally crossed with three *Cas9* lines driven by *nanos* (*nos*), *vasa* (*vas*), and *Ubiquitin-63E* (*Ubi*) promoters and were sufficient to ensure complete penetrance of both female lethality / masculinization and male sterility in each reciprocal cross (Fig. S1, S2). Gender frequencies and fertility in trans-heterozygotes were compared to those in corresponding progeny of control crosses with *Cas9* (bar groups to the left, solid lines) or *dgRNAs* (top panels, dashed lines) and *wt* flies. Each bar shows an average gender frequency and one standard deviation. Statistical significance was calculated with a *t* tests assuming unequal variance, and for male sterilization, *P* values were calculated using Pearson’s Chi-squared test for contingency tables (red *). (*P* > 0.01**, *P >* 0.001***). (**B**) Order of targeted gene in sex-determination pathway (top) and corresponding knockout phenotype in progeny. Phenotypes of *dgRNAs* directed-knockouts and intersex morphology in comparison to *wt* ♀ and ♂. *βTub, Sxl* knockouts ♀ perish during pupal stages (Fig. S3). *dgRNA^βTubJra^/+; nos-Cas9/*+ intersexes (⚥), but not *dgRNA^βTub,DsxF^/+; nos-Cas9/*+ ⚥, had sex combs, magnified inside inserts. (**C**) Variable expressivity of the number of sex comb bristles in *βTub, Tra* knockouts ⚥. (**D**) Internal reproductive organs in *wt* females: two ovaries (ov), seminal receptacle (*sr*), double spermatheca (*sp*), two accessory glands (*ag*), and uterus (ut). (**E**) Many *dgRNA^βTubJra^/+; nos-Cas9/*+ ⚥ had one rudimentary ovary, and organs that resembled male accessory glands. (**F**) Many *dgRNA^βTub,DsxF^/+; nos-Cas9/*+ ⚥ developed only a single ovary often times not connected with an oviduct and had organs that resembled male-specific accessory glands. (**G-H**) Male internal reproductive system in *dgRNA^βTub,Sxl^/+; nos-Cas9/*+ ⚥. In comparison to *wt* testis (**I**), elongated cysts with maturing spermatids were not found in the *dgRNA^βTub,Sxl^* testis (**H, I**: arrows).

In terms of phenotypes, we found that the 100% of the *dgRNA^βTub,Sxl^* knockout females perished during pre-adult stages with the majority dying during a pupal transition (Fig. S3). For intersex phenotypes, fertility was always compromised, however variable expressivity was observed as the extent of anatomical masculinization varied between individuals and was more pronounced in the *dgRNA^βTub,Tra^* knockouts as compared to the *dgRNA^βTub-DsxF^* (Fig. 2B, Table S4). For example, *dgRNA^βTub,Tra^* knockout intersexes had sexcombs with variable bristle numbers (Fig. 2B–C, Table S4), and rarely developed more than one rudimentary ovary (Fig. 2D–E, Table S4). Moreover, molecularly the *dgRNA^βTub,Tra^* knockout intersexes expressed both female and male-specific alternative splice variants of *dsx* gene (Fig. S4), presumably due to the absence of Tra which is important for inhibiting the male-specific and promoting the female-specific alternative splicing of *dsx ^37^*. In contrast, the *dgRNA^βTub,DsxF^* knockout intersexes were not observed to develop sexcombs, and some instersexes had normal ovaries enabling them to become gravid, although unable to oviposit (Fig. 2B, F, Table S4).

In regard to male infertility phenotypes, to visualize the anatomy of testes and developing spermatids in the F_1_ sterile males, we generated a transgenic line expressing eGFP under control from the *βTub85D-promoter* (*βTub-GFP*) to fluorescently label the testes and sperm (Fig. S1), and introgressed it with the *dgRNA* strains. When crossed with homozygous *nos-Cas9*, the trans-heterozygous *dgRNA^βTub,Sxl^/+; βTub-GFPInos-Cas9* F_1_ sterile males had fully developed coiled testes and accessory glands (Fig. 2G), however spermatid development was completely disrupted with phenotypes consistent with previous *βTub* disruption reports ^29^. For example, only round cysts and early spermatocytes were identified in the testes of sterile males marked with GFP (Fig. 2H), while *wt* testes had robust GFP-labeled cysts with elongated late spermatids (Fig. 2I). Moreover, given that the *βTub-GFP* labels testes/sperm, this tool enabled us to explore the internal anatomy of reproductive systems in intersexs to search for putative male testes-like structures. Although no GFP-positive testes were identified in either *dgRNA^βTub,Tra^ or gRNA^βTub,DsxF^* knockout intersex individuals (n>20, Table S4) paired putative male accessory gland like organs were present in both intersex types (Fig. 2D, E-G). Finally, to confirm the molecular changes that resulted in knockout phenotypes, we sequenced both targeted loci from individual F_1_ flies. As expected, compared to the control flies (n=32) each examined double knockout fly (n=20) had mosaic indels precisely at the cleavage sites that prevented sequencing through both ends of PCR amplicons (Fig. S5, S6, Table S2).

### Complete Penetrance Resulting From Zygotic Expression

Maternal deposition of Cas9/gRNA complexes into developing embryos is sufficient to ensure non-Mendelian inheritance of mutations in receiving progeny, even if those progeny do not genetically inherit the genes encoding the editing components, and this phenomenon is known as dominant maternal effect ^38^. To extend this work, we aimed to test whether paternal inheritance of one of the core components (i.e. *Cas9* or *dgRNA*), combined with maternal deposition of the compatible component, would be sufficient to generate heritable mutations. For the first combination, matings between homozygous Cas9 fathers and heterozygous dgRNA expressing mothers were not sufficient to induce mutations (n=12), or knockout phenotypes (n=252, N=6), in F_1_ progeny that did not inherit the dgRNAs as a gene, presumably a result of a short dgRNA half-life in the absence of Cas9 during maternal deposition (Fig. 3B, Table S2). Moreover, matings between heterozygous Cas9 fathers and homozygous dgRNA-expressing mothers resulted in male sterility and female lethality/masculinization phenotypes in all trans-heterozygous F_1_ progeny that inherited the *Cas9* gene (n=2191,N=27), while all F_1_ progeny that inherited only the dgRNA-encoding genes maintained normal features (n=2640, N=27)(Fig. 3A, Table S7). Moreover, crosses between heterozygous Cas9 mothers and homozygous dgRNA-expressing fathers resulted in male sterility and female lethality/masculinization phenotypes in all trans-heterozygous F_1_ progeny (n=3019, N=36)(Fig. 3A, Table S7). Additionally, maternal contribution of Cas9 protein was sufficient to induce intersex phenotypes in progeny that did not receive the *Cas9* gene when targeting *tra* or *dsx* (n=782, N=24), demonstrating the dominant maternal effect (Fig. 3A). However, maternal contribution of Cas9 only by *Ubi-Cas9* (n=0 (# surviving females), N=4), but not *nos-Cas9* nor *vas-Cas9* (n=556, N=8), induced *dgRNA^βTub,sd^/+; +/+* female lethality indicating that promoter strength likely plays an important role in mutation efficiency (Fig. 3A-B). Interestingly, despite the lack of lethality phenotypes in females receiving Cas9 protein maternally loaded from *nos-Cas9* and receiving the *dgRNA^βTub,Sxl^* gene, these surviving females had mosaic indels at the *Sxl* locus (n=2, Table S2, Fig. 3B). Similarly, all male progeny that inherited only the *dgRNA* genes (n=1490, N=36), and had maternally loaded Cas9 protein, were fertile regardless of Cas9 strain used (Fig. 3A), though each genotyped male (n=6) had mosaic indels at the *βTub* locus (Table S2, Fig. 3B). Taken together, these results indicate that paternal inheritance of gRNAs along with maternal deposition of Cas9 into developing embryos, in the absence of Cas9 inherited as a gene, was sufficient to induce detectable biallelic mosaicism, although penetrance was incomplete depending on gene targeted.

**Fig. 3.**
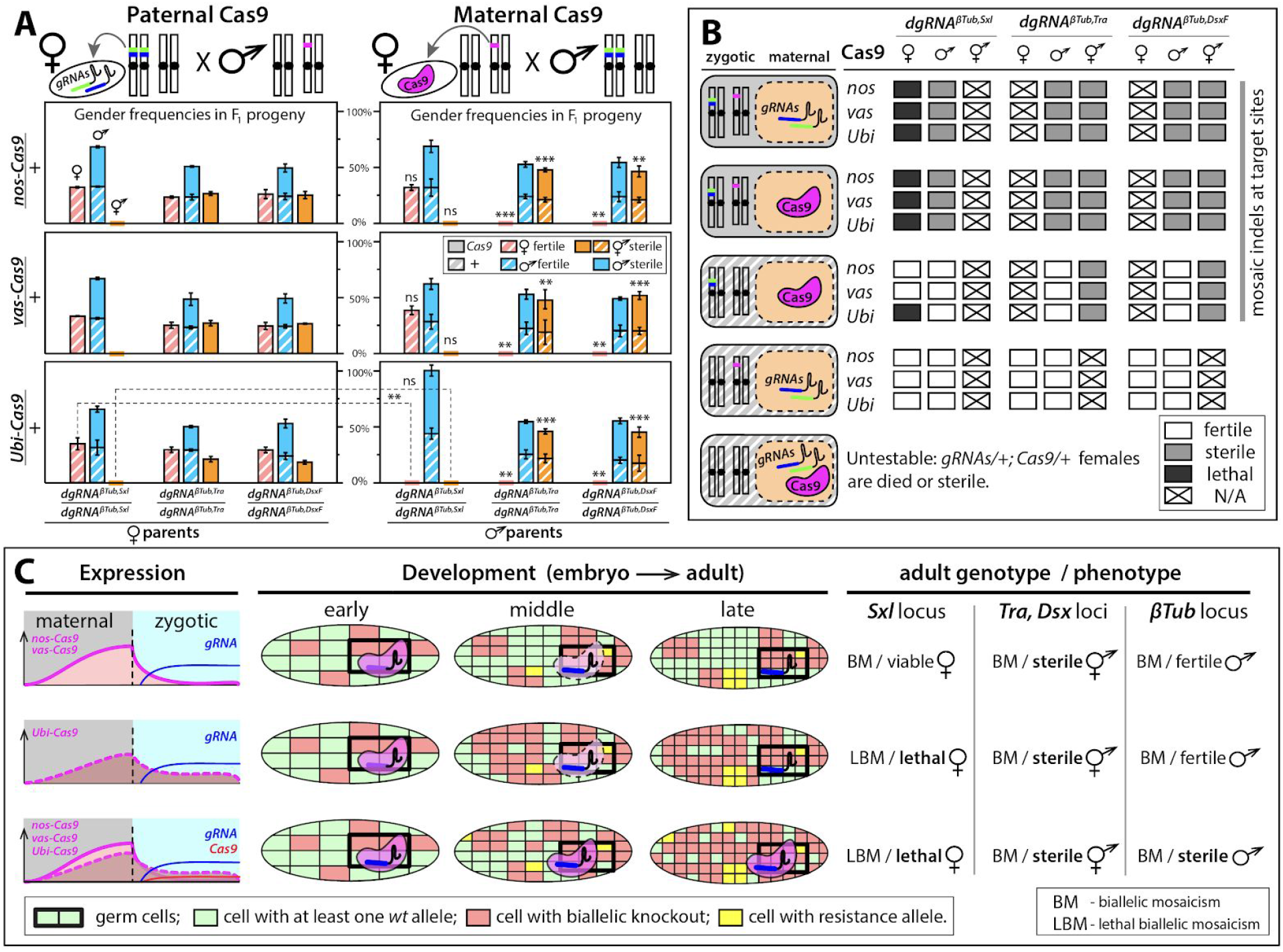
Zygotic expression of CRISPR components ensures 100% penetrance. (**A**) Genetic quantification of dominant effect by maternal loading of Cas9. Genotypes, gender frequencies, and fertility of flies generated by reciprocal crosses between homozygous *dgRNAs* and heterozygous *Cas9* flies. Progeny from crosses with heterozygous paternal *Cas9* (left panels) and heterozygous maternal *Cas9* (right panels). Each bar shows an average gender frequency and one standard deviation. Statistical significance was calculated with *t* tests assuming unequal variance. (*P* > 0.01**, *P >* 0.001***). Striped bars indicate inheritance of *Cas9* as a gene, while solid bars indicate inheritance of + allele. (**B**) Combinations of genotypes and maternal/zygotic contributions in embryos, and their penetrance. (**C**) Accumulation of high levels of biallelic mosaicism (BM) throughout development leads to the loss of gene function at the organismic level and ensures complete penetrance of induced phenotypes: lethality (lethal biallelic mosaicism (LBM)), female masculinization, or male sterility. Complementation of gene function in some cells by uncleaved *wt* alleles, or resistance alleles generated by NHEJ, is not sufficient to rescue the induced phenotype at the organismic level and therefore 100% of trans-heterozygous progeny have the induced phenotypes. Boxes get smaller and more abundant as cells divide.

### pgSIT Males Sexually Compete for Mates

Given the simplicity and consistency of generating sterile males (Fig. 1A), pgSIT could potentially be used in the future to mass produce and release eggs into the environment to suppress target populations. A potential application of pgSIT would be the suppression of populations of *Ae. aegypti*, the mosquito vector of dengue, Zika and Chikungunya. To explore how the pgSIT approach may fare against currently-available self-limiting suppression technologies - namely RIDL, fsRIDL and IIT - we simulated release schemes for each of these technologies using the MGDrivE simulation framework ^39^. This framework models the egg, larval, pupal and adult mosquito life stages with overlapping generations, larval mortality increasing with larval density, and a mating structure in which females retain the genetic material of the adult male with whom they mate for the duration of their adult lifespan ^39^. We consider releases into a randomly-mixing population consisting of 10,000 adult females, with model and intervention parameters listed in Table S9. To parameterize the mating competitiveness of pgSIT males, we performed a mating competition assay (Fig. 4A). From this experiment, we discovered that pgSIT-generated males were able to court, mate and successfully compete with *wt* males. A reduced egg hatch rate of 47.9%±13.8% for 1 *wt* and 1 pgSIT male vs. 85.1%±13.5% for 2 *wt* males (N=5, *P*>0.003) (Fig. 4B) was consistent with a mating competitiveness of pgSIT males of 78% that of *wt* males.

**Fig. 4.**
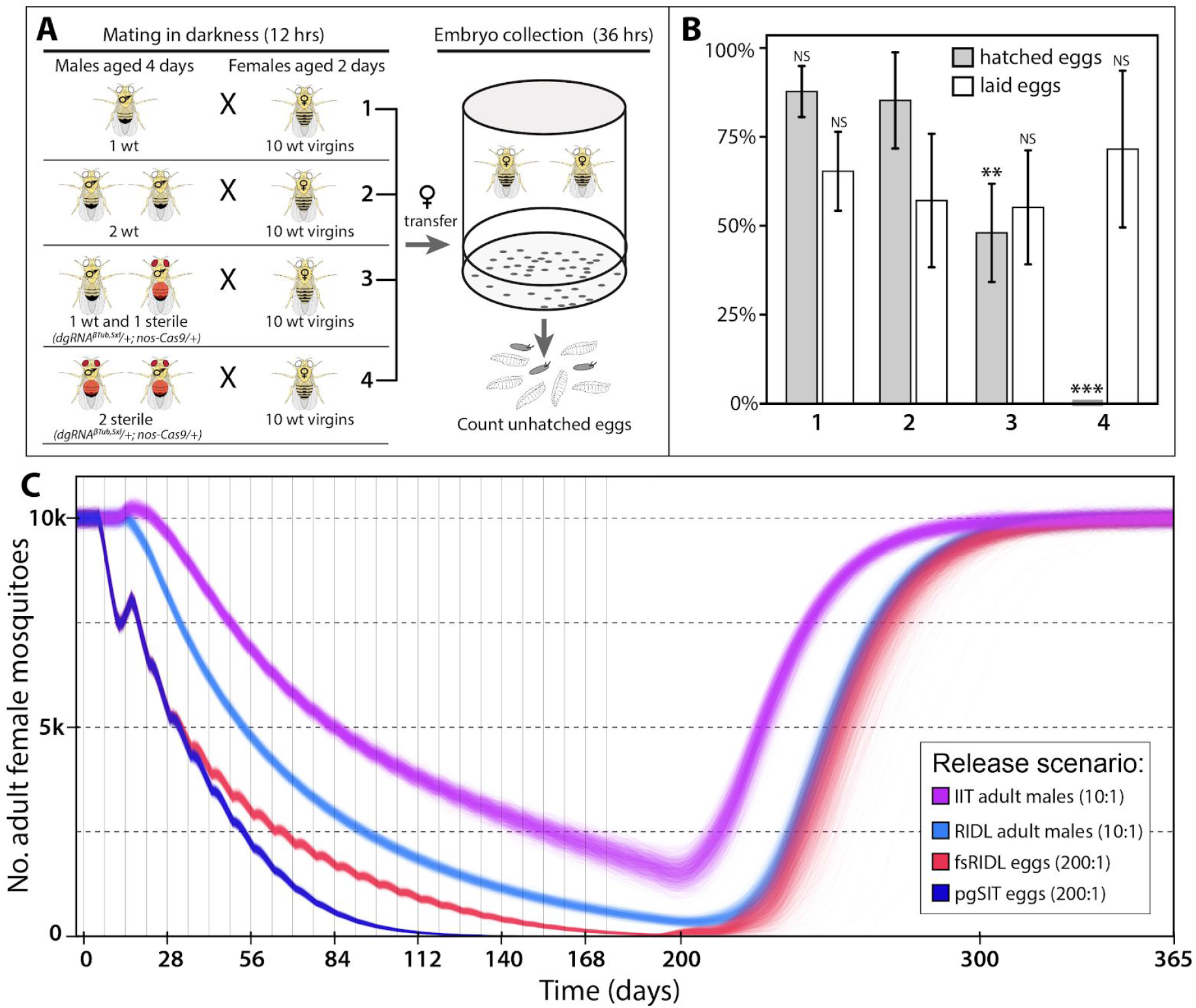
Competitiveness of pgSIT males and modeling data. (**A**) An experimental setup to estimate the mating competitiveness of *dgRNA^βTub,Sxl^/+; nos-Cas9/+* sterile males (marked with red) competing against *wt* males to secure matings with *wt* females. A mated female is resistant to the next mating for around 24 hours ^57,60^, and the mating success of sterile males was evaluated by fertility decrease (aka. increase of unhatched egg rate). (**B**) Bars graphs percentages of laid and hatched eggs. Numbers of laid eggs were normalized against the highest egg number (n=199) to convert them to the percentile (Table S8). The presence of one sterile male resulted in a significant decrease in female fertility (#3 vs #2) that could not be accounted by removal of one *wt* male (#2 vs #1). Statistical significance was calculated with a *t* test assuming unequal variance comparing group #3 to #2 (*P* > 0.003**, *P* > 0.0001***). (C) Model-predicted impact of releases of pgSIT eggs on *Aedes aegypti* mosquito population density with comparison to releases of *Wolbachia*-based incompatible insect technique (IIT), RIDL (release of insects carrying a dominant lethal gene) and fsRIDL (female-specific RIDL). Releases are carried out weekly over a six month period with release ratios (relative to wild adults) shown in the key. Model predictions were computed using 2000 realizations of the stochastic implementation of the MGDrivE simulation framework ^61^ for a randomly-mixing *Ae. aegypti* population of 10,000 adult females and model parameters described in Table S9. Notably, pgSIT releases outcompete those of all other suppression technologies, showing the highest potential to eliminate the local population.

We simulated weekly releases of adult males for RIDL and IIT and eggs for fsRIDL and pgSIT over a 6 month period (Fig. 4C). Adult release ratios were 10 adult RIDL/IIT males per wild adult, following the precedent of a field trial of *Ae. aegypti* RIDL mosquitoes in Brazil ^40^, and egg release ratios were 200 eggs per wild adult, given that female *Ae. aegypti* produce ~20 eggs per day in temperate climates ^41^. Results from these simulations suggest that systems for which eggs are released (pgSIT and fsRIDL) result in the most rapid population suppression in the first three weeks as released eggs quickly hatch as larvae and reduce the survival of fertile larvae as a consequence of density-dependent larval competition. The pgSIT approach shows the greatest suppression from the end of the first month on, and the greatest potential to eliminate the population during the release period. This is due to the higher mating competitiveness of pgSIT males (78% that of *wt* males) c.f. fsRIDL males (~5% that of *wt* males, based on RIDL field trials in the Cayman Islands ^42^ and Brazil ^40^), which becomes a dominant factor at low population densities when greater consumption of larval resources by released immature forms has less impact on suppression. Population suppression resulting from 10:1 releases of adult RIDL males trails that for releases of fsRIDL eggs by 2-3 weeks due to the delay in impact on density-dependent larval competition; but is similar in magnitude. Equivalent releases of adult IIT males are less impactful for the strategy we consider, in which male incompatibility is induced through *Wolbachia* infection and the chance of an unintended release of *Wolbachia*-infected females interfering with suppression is reduced through low-level irradiation ^43,44^, resulting in the longevity of released IIT males being roughly halved ^45^.

In sum, these results suggest that pgSIT has greater potential to eliminate local *Ae. aegypti* populations than currently-available population suppression technologies. The results also appear highly robust to variation in the lifespan and mating competitiveness of pgSIT adult males (Fig. S7). For weekly releases of 200 pgSIT eggs per wild adult, simulations suggest a wide range of parameter values for which local *Ae. aegypti* elimination could be reliably achieved (male mating competitiveness > ~25%, lifespan reduction < ~75%). Elimination could also be reliably achieved for smaller releases of 100 pgSIT eggs per wild adult (male mating competitiveness > ~50%, lifespan reduction < ~50%).

## Discussion

### Genetic Variation and Resistance Unlikely to Hinder PgSIT

CRISPR has empowered us to develop a novel system (pgSIT) to enable the release of eggs to ensure all progeny surviving to adulthood are sterile males - a feat never before possible. This is accomplished by using advanced molecular genetics to simultaneously sterilize males and eliminate females. Importantly, pgSIT relies exclusively on highly efficient CRISPR-mediated DNA cleavage and NHEJ-based repair and does not rely on HDR. Therefore, generation of resistance alleles that can curtail CRISPR-mediated gene drives ^10^ does not limit efficacy of pgSIT, as absolute disruption of *wt* alleles is not required to ensure complete penetrance of the induced phenotype when targeting essential genes. Additionally, accumulation of resistance is unlikely to pose an issue for pgSIT since homozygous strains are raised separately then mated to produce sterile males which do not generate viable progeny, limiting the selection pressure on the target sites. Given that the role of pgSIT males is simply to seek out wild females, mate, and thereby reduce fecundity, natural genetic diversity in the wild is also not likely to pose a problem. These considerations result in the extraordinary efficiency and robustness of pgSIT to be used directly for population control.

### Why is PgSIT 100% Efficient?

In terms of underlying mechanism for pgSITs extreme efficiency, we determined that zygotic activity of the Cas9/gRNA complexes ensures continuous biallelic mosaicism of targeted alleles throughout development resulting in complete penetrance of desired phenotypes, although variable expressivity may still be observed depending on the gene targeted and on the timing and strength of Cas9 promoter (Fig. 3C). Moreover, paternal inheritance of dgRNAs along with maternal deposition of Cas9 into developing embryos, in the absence of Cas9 inherited as a gene, was also sufficient to induce detectable biallelic mosaicism for all genes targeted (*βTub, dsx, tra, sxl*), although penetrance was incomplete depending on gene targeted. For example, maternal deposition of Cas9 alone was sufficient to induce intersex phenotypes (*dsx, tra*), however it was insufficient to phenotypically ensure male sterility (*βTub*), and depended on the strength of the promoter maternally depositing Cas9 to ensure female death (*sxl*) via lethal biallelic mosaicism (Fig. 3B–C). Taken together, these observations suggest that rates of biallelic mosaicism which ensure complete penetrance depend exclusively on whether components (i.e. Cas9 and gRNA) are inherited as genes or maternally deposited. Additionally, regardless of how the components are inherited, if rates of biallelic mosaicism are over a critical threshold, which is specific to each gene targeted, then complete penetrance can be achieved. Mechanistically, this technology demonstrates a fundamental advance in genetics by which somatic biallelic disruptions in essential genes, that previously conferred recessive phenotypes, get simultaneously converted by pgSIT in many somatic cells resulting in dominant fully penetrant phenotypes in a single generation.

### Steps Toward Developing pgSIT in Disease Vectors

The simplicity of system provides a rationale for developing pgSIT in many insect species including disease vectors and agricultural pests. Importantly the technology does not rely on chromosome translocations, chemosterilants, irradiation, antibiotics or bacterial infections, which can severely compromise the fitness and mating competitiveness of released sterile males. To implement pgSIT in disease vectors, many genes important for female viability and male fertility can be targeted. For example, given the functional conservation, *dsx* ^46,47^ and *βTub* ^48,49^ could be tested initially in mosquitoes, but there are plenty of other female/male specific genes that could be targeted ^50,51^. Notwithstanding, while there are many genes to target, care must be taken in target gene selection to ensure minimal negative impacts on male fitness and courtship behavior. Moreover, given that highly efficient, genomically-encoded, Cas9-expressing strains that have already been developed in major dengue and malaria disease vectors including *Ae. aegypti* ^36^, *Anopheles gambiae* ^5^, and *Anopheles stephensi* ^4^, suggests pgSIT may be trivial to develop in these species. To efficiently utilize pgSIT for mosquitoes, we envision the development of a rearing facility to propagate homozygous Cas9 and dgRNA expressing strains separately. An automated workflow would also need to be implemented to sex-sort immature stages (e.g. Cas9 females with dgRNA males) and combine into cages for maturation, mating and propagation of eggs. Sex sorting can be achieved in a number of ways including mechanical size separation, automated copas sex sorting platform (Union Biometrica) combined with a genetic sexing strain, or automated robotic optical sorting and therefore should not be an insurmountable limitation (reviewed in ^52,53^). It should be noted that pgSIT would be quite effective for insect species by which eggs could be stored desiccated in dipause for long periods, for example, *Ae. aegypti* and *Ae. albopictus*, to enable scalable egg accumulation for inundative releases. An efficient pgSIT egg production facility, could distribute pgSIT eggs to remote field sites all over the world via drones, by which they could simply be hatched, reared, and released, eliminating the logistical burden of manual sex-sorting, sterilization, and releasing fragile adult males in the field, thereby increasing scalability, and efficiency, enabling broader wide-scale population suppression capacity (Fig. S8).

### Potential to Eliminate Disease Vector Populations

Mathematical modeling of pgSIT alongside currently-available self-limiting suppression technologies – RIDL, fsRIDL and IIT – suggests that pgSIT has the highest potential to eliminate local *Ae. aegypti* populations and highlights the relative strengths of the pgSIT approach, even before the cost-effectiveness and scalability of egg releases are taken into account (Fig. 4C). Egg releases result in rapid population suppression from the outset, as hatching larvae consume resources that would otherwise be available to fertile larvae. A beneficial property shared by both pgSIT and fsRIDL is that all released eggs can result in hatching larvae, as female lethality occurs after the larval stage, resulting in maximum consumption of larval resources by released immature forms. We predict pgSIT to achieve greater suppression than fsRIDL and RIDL, in their current forms, due to the substantially higher mating competitiveness of pgSIT males (~78% that of *wt* males) c.f. RIDL males (~5% that of *wt* males). Mating competitiveness is a dominant factor in achieving local elimination, as once initial suppression has been achieved, larval resources are abundant and hence greater consumption by released immature forms is less impactful. Improving the mating competitiveness of RIDL males is conceivably an engineering problem hinging on reducing toxin leakage following rearing with tetracycline; however the cause of such a large reduction in mating competitiveness is, to our knowledge, unclear. Regardless, pgSIT has an additional advantage over fsRIDL when it is preferred that introduced transgenes do not persist in the environment for more than a generation following their final release. Additional excitement for pgSIT stems from its potential to eliminate local *Ae. aegypti* populations for a wide range of lifespan and mating competitiveness parameter values (Fig. S7C), suggesting some wiggle room when porting to other species. Simulations also suggest that pgSIT may be capable of eliminating local populations given smaller release ratios (Fig. S7D-F). Combined with the feasibility and cost-effectiveness of mass rearing and release of pgSIT eggs, this points to a highly promising technology for the suppression of local populations of insect agricultural pests and disease vectors.

## Materials and Methods

### CRISPR target site design

To confer female lethality and male sterility, target sites for guide RNAs (gRNAs) were chosen inside female-specific exons of sex-determination genes, *Sex Lethal* (*Sxl*), *Transformer* (*tra*), and *Doublesex* (*dsx*), and in male specific genes, *βTubulin 85D (βTub), fuzzy onions (fzo), Protamine A* (*ProA*), and *spermatocyte arrest* (*sa*), respectively. CHOPCHOP v2 ^54^ was used for choosing gRNA target sites from specified sequence in *Drosophila* genome (dm6) to minimize the off-target cleavage. Due to the alternative splicing, functional Sxl and Tra proteins are produced only in *Drosophila* females ^26,27^, while two versions of Dsx protein - female (Dsx^F^) or male (Dsx^M^) - are made each in the corresponding gender ^28^ (Fig. 1B). The gRNA target for *βTub* was chosen in the vicinity to the *βTub85D^D^* (*B2t^D^*) mutant allele ^29^. Sequences of gRNA target sites are presented in Fig. S1.

### Design and assembly of constructs

Gibson enzymatic assembly method was used to build all constructs ^55^. The previously described plasmid harboring the *SpCas9-T2A-GFP* with nuclear localization signals (NLS) flanking SpCas9 coding sequence and the Opie2-dsRed transformation marker was used to build *Drosophila* Cas9 constructs. This plasmid was used for *Ae. aegypti* transgenesis and had both piggyBac and an attB-docking sites (addgene #100608)^36^. The *Ae. aegypti* promoter was removed from the plasmid by cutting at NotI & XhoI sites and replacing it with *Nanos* (*nos*), or *Ubiquitin-63E* (*Ubi*), or *Vasa* (*vas*) promoter (Fig. S1). Promoter fragments were PCR amplified from *Drosophila* genomic DNA using the following primers: nos-F, nos-R, Ubi-F, Ubi-R, vas-F, and vas-F (Table S10). To generate constructs with a single gRNA, *Drosophila* U6-3 promoter and guide RNA with a target, scaffold, and terminator signal (gRNA) was cloned at the multiple cloning site (MCS) between the *white* gene and an attB-docking site inside a plasmid used for *D. melanogaster* transformation ^56^. For the first plasmid in this series, U6-3-gRNA^βTub^, *Drosophila* U6-3 promoter was amplified from *Drosophila* genomic DNA with U6-1F and U6-2R primers while the complete gRNA was PCR-assembled from two ultramer^®^ gRNA-3F and gRNA-4R oligos synthesized by Integrated DNA Technology (IDT). To improve the efficiency of termination of gRNA transcription, a termination signal with 11 thymines was used in our design. In the successive plasmids, the U6-3 promoter and gRNA’s scaffold was amplified from the U6-3-gRNA^βTub^ plasmid using the overlapping middle oligos designed to replace 20 bases that constitute a gRNA target (U6-1AF, U6-2A/B/CR, gRNA-3A/B/CF, and gRNA-4AR), and replaced by digesting the same plasmid at AscI and NotI sites. To assemble the set of plasmids with double gRNAs (dsRNAs), the U6-3 promoter and gRNA was amplified as one fragment from the single gRNA (sgRNA) plasmids targeting female sex-determination genes with 2XgRNA-5F and 2XgRNA-6R primers, and cloned inside the U6-3-gRNA^βTub^ plasmid that was linearized at a BamHI site between the *white* gene and the U6-3 promoter. Each dgRNA plasmid had the same gRNA^βTub^ targeting *βTub85D* and a different gRNA targeting *Sxl, tra*, or *dsxF* expressed independently in the same direction (Fig. S1). *Drosophila* Cas9 plasmids and gRNA plasmids generated for this study were deposited at Addgene (Fig. S1). To build the *βTub85D-GFP* construct, a 481bp fragment directly upstream of *βTub* coding sequence was PCR amplified from *Drosophila* genomic DNA with βTub-F and βTub-R primers and cloned upstream of *GFP* into the *white* attB-docking site plasmid described above.

### Fly genetics and imaging

Flies were maintained under standard conditions at 25 °C. Embryo injections were carried at Rainbow Transgenic Flies, Inc. (http://www.rainbowgene.com) The *Cas9* and gRNA constructs were inserted at the PBac{y+-attP-3B}KV00033 on the 3^rd^ chromosome (Bloomington #9750) and the P{CaryP}attP1 site on the 2^nd^ chromosome (Bloomington #9750), respectively; while *βTub-GFP* construct was inserted at the M{3XP3-RFP.attP’}ZH-86Fa on the 3^rd^ chromosome (Bloomington #24486) (Fig S1). Transgenic flies were balanced with w^1118^; CyO/sna^Sco^ and w^1118^ and TM3, Sb^1^/TM6B, Tb^1^; and double balanced with w^1118^; CyO/Sp; Dr^1^/TM6C,Sb,Tb^1^. The *βTub-GFP* (on the 3rd chromosome) was double balanced and introgressed with *gRNA^βTub,Sxl^*, *gRNA^βTub,Tra^*, and *gRNA^βTub-DsxF^*, each on the 2nd chromosome, to generate trans-heterozygous balanced stocks (*dgRNA/CyO; βTub-GFP/TM6C,Sb,Tb*).

To test the efficiency of knockouts and corresponding phenotypes caused by sgRNAs, seven flies of each gender were crossed to generate trans-heterozygous F_1_ *sgRNA/+; nos-Cas9/+* flies for each combination of sgRNA; and their external morphology and fertility were examined. Both transgenes were identified on a fluorescent stereo microscope with w+ eyes (*sgRNA, dgRNA*) and dsRed (*Cas9*). The sgRNA lines that caused knockout phenotypes were further tested as homozygous stocks with *nos-Cas9* flies in both directions using 10+ flies of each gender. DgRNAs lines were tested bidirectionally with homozygous *nos-Cas9, vas-Cas9*, and *Ubi-Cas9* lines. In addition, *sgRNA, dgRNA* and *Cas9* homozygous lines were crossed to w-flies in both directions to provide the comparison control. To test for the non-Mendelian dominant maternal effect of Cas9 loaded as protein into embryos ^38^, homozygous *dgRNA* flies were crossed to heterozygous *Cas9* flies; and phenotypes of *dgRNA/+; +/TM3, Sb* progeny with either maternal Cas9 or paternal Cas9 were compared. The F_1_ progeny from crosses with the paternal Cas9 served as a control group to examine the dominant maternal effect of Cas9. To test fertility of generated knockout flies with and without the Cas9 gene, batches of 10-20 F_1_ males and females, or intersexes, were crossed to 15-20 female virgin and male flies, correspondingly, from w- and/or Cantos S stock lines. Three or four days after the cross, the flies were passaged into fresh vials, and in a week, both vials were examined for presence of any viable progeny. The fertility of an entire batch was scored as 100% when viable larvae were identified in a vial, or 0% when no progeny hatched in both vials. The vials containing intersexes and *wt* males were also examined for presence of laid eggs. All crosses were repeated at the minimum three times to generate means and standard deviations for statistical comparisons and thus measure consistency and robustness of the results.

Flies were scored, examined, and imaged on the Leica M165FC fluorescent stereo microscope equipped with the Leica DMC2900 camera. To generate images of adult flies, image stacks collected at different focal plates were compiled into single images in Helios Focus 6, which were edited in Adobe Photoshop CS6. To study internal anatomical features of intersex flies and sterile males, their reproductive organs were dissected in PBS buffer, examined, and imaged. To estimate the variation of knockout phenotypes, around 10-20 flies were dissected for each tested genotype.

### Developmental stage of *Sxl* lethality

To identify the developmental stage at which *Sxl* knockout females die, egg hatching and larval death rates were quantified for the *dgRNA^βTub,Sxl^/+; nos-Cas9/+* trans-heterozygous flies. To quantify the egg hatching rate, three replicate crosses, each with 20-30 homozygous *nos-Cas9* female virgins and 10-20 *dgRNA^βTub,Sxl^* males, were set up in embryo collection cages (Genesee Scientific 59-100) with grape juice agar plates. Three embryo collection cages with w-flies served as a comparison control. Batches of around 200 laid eggs were counted from each collection cage and followed for over 36 hours to count the number of unhatched eggs. To quantify the rate of larval death, two batches of 50 emerged larvae were transferred from each agar plate to separate fly vials with food and raised to adults, then a number and sex of emerged adults were recorded. To quantify the lethality at a pupal stage, a number of dead pupae was also recorded for each vial.

### RT-PCR of female- and male-specific splice transcripts of *Dsx*

To assess the effect of *tra* knockout on *dsx* splicing, we screened for female- and male-specific mRNA of *dsx* in *tra* knockout intersexes. Total RNA were extracted from adult w-male, w-female, and *tra* knockout (*dgRNA^βTub,Tra^/+; nos-Cas9/+*) intersex flies following the standard protocol of the MirVana miRNA isolation kit (Ambion). To remove DNA contamination, 2 μg was treated with TURBO^™^ DNase using the TURBO DNA-free^™^ Kit (Ambion). *Dsx* female and male splice variants were amplified with the Superscript ^®^ III One-Step RT-PCR Kit (Invitrogen) following the protocol. The same forward primer, Dsx-RT-1F, and two different reverse primers, DsxF-RT-2R and DsxM-RT-3R, were used to amplify either female or male transcripts, respectively. 10 μL of PCR products were run on a 1% agarose gel to test PCR specificity, and the remaining 40 μL were purified using a QIAquick PCR purification kit (QIAGEN) or, when double bands were identified on a gel, gel-purified with a Zymoclean ^™^ Gel DNA Recovery Kit (Zymo Research), then clean amplicons were sequenced in both directions using Sanger method at Source BioScience (https://www.sourcebioscience.com).

### Genotyping loci targeted with gRNAs

To examine the molecular changes that caused female lethality or masculinization and male sterility in the flies carrying Cas9 and gRNAs, four genomic loci that include targets sites for four functional gRNAs (Fig. S1) were amplified and sequenced. Single-fly genomic DNA preps were prepared by homogenizing a fly in 30μl of a freshly prepared squishing buffer (10mM Tris-Cl pH 8.0, 1mM EDTA, 25mM NaCL, 200 μ/mL Proteinase K), incubating at 37°C for 35 minutes, and heating at 95°C for 2 minutes. 2 μl of genomic DNA was used as template in a 40 μL PCR reaction with LongAmp^®^ Taq DNA Polymerase (NEB). The following primers (Table S10) were used to amplify the loci with the corresponding gRNA targets: βTub-1AF and βTub-2AR for *βTubulin 85D;* Sxl-3BF and Sxl-4AR for *Sex lethal;* Tra-5F and Tra-6R for *Transformer*, Dsx-7F and Dsx-8R for *Double sex*. PCR products were purified using a QIAquick PCR purification kit (QIAGEN), and sequenced in both directions with Sanger method at Source BioScience. To characterize molecular changes at the targeted sites, sequence AB1 files were aligned against the corresponding reference sequences in SnapGene^®^ 4 and / or Sequencher™ 5.

### Competition assay of sterile males

To evaluate the competitiveness of the *βTub* knockout (*gRNA^βTub,Sxl^/+; nos-Cas9/*+) males, their ability to secure matings with females in the presence of *wt* males was evaluated. The w-males share the same genetic background with the *βTub* knockout males, and provide an ideal comparison. Two *wt*, one *wt*, one *wt* plus one *βTub* knockout, or two *βTub* knockout males were placed into a fly vial with ten w-virgins isolated on yeast paste for two days and allowed to court and mate with the females overnight (12 hours) in the dark. To increase the male courtship drive, freshly emerged *dgRNA^βTub,Sxl^/+ ; nos-Cas9/+* and *wt* males were isolated from females and aged for four days before the competition assay. *Drosophila* females mate with multiple males during a lifespan; and in the absence of sperm transferred during copulation, female abstinence lasts for one day postcopulation (Peng et al. 2005). Therefore, after 12 hours of mating, all males were removed from the vials while the females were transferred into small embryo collection cages (Genesee Scientific 59-100) with grape juice agar plates. Three batches of eggs were collected within 36 hours and unhatched eggs were counted. The decrease in fertility, estimated by a number of unhatched eggs, indicated the ability of a *gRNA^βTub,Sxl^/+; nos-Cas9/+* male to score successful matings with females in the presence of a *wt* male; and thus provided a readout of the competitiveness of *βTub* knockout males. A single *wt* male was used to test its ability to inseminate each of ten females in 12 hours, and thus discriminate between a true competition or a dilution effect of two *wt* males.

### Mathematical modelling

To model the expected performance of pgSIT at suppressing local *Ae. aegypti* populations in comparison to currently-available self-limiting suppression technologies - RIDL, fsRIDL and IIT - we simulated release schemes for each using the MGDrivE simulation framework ^39^ (https://marshalllab.githubio/MGDrivE/) This framework models the egg, larval, pupal and adult mosquito life stages (both male and female adults are modeled) implementing a daily time step, overlapping generations and a mating structure in which adult males mate throughout their lifetime, while adult females mate once upon emergence, retaining the genetic material of the adult male with whom they mate for the duration of their adult lifespan. Density-independent mortality rates for the juvenile life stages are assumed to be identical and are chosen for consistency with the population growth rate in the absence of density-dependent mortality. Additional density-dependent mortality occurs at the larval stage, the form of which is taken from Deredec *et al*. (2011)^58^. The inheritance patterns for the pgSIT, RIDL, fsRIDL and IIT systems are modeled within the inheritance module of the MGDrivE framework ^39^, along with their impacts on adult lifespan, male mating competitiveness and pupatory success. We implement the stochastic version of the MGDrivE framework to capture the random effects at low population sizes and the potential for population elimination. We simulated weekly releases over a period of 6 months into a randomly-mixing population consisting of 10,000 adult females at equilibrium, with *Ae. aegypti* life history and intervention parameter values listed in Table S9.

### Statistical analysis

Statistical analysis was performed in JMP 8.0.2 by SAS Institute Inc. Three to five biological replicates were used to generate statistical means for comparisons. *P* values were calculated for a two-sample Student’s *t*-test with unequal variance. To test for significance of male sterilization, Pearson’s Chi-squared tests for contingency tables were used to calculate *P* values.

### Data Accessibility

Complete annotated plasmid sequences and plasmid DNA are publically available for order at Addgene. Transgenic flies have been made available for order from Bloomington *Drosophila* stock center (see Figure S1 for details).

## Acknowledgements

This work was supported in part by funding from NIH grants including NIH-K22 Career Transition award (5K22AI113060), an NIH Exploratory/Developmental Research Grant Award (1R21AI123937) awarded to O.S.A, and a Defense Advanced Research Project Agency (DARPA) Safe Genes Program Grant (HR0011-17-2-0047) awarded to O.S.A. and subcontracted to J.M.M. We thank Ethan Bier (UCSD), Anthony A. James (UCI), and Craig Montell (UCSB) for their insightful discussions, comments and edits.

## Author Contributions

O.S.A and N.P.K. conceived and designed experiments. N.P.K. and J.L. performed molecular and genetic experiments. H.M.S.C., S.L.W. and J.M.M. conducted the mathematical modeling. All authors contributed to the writing, analyzed the data, and approved the final manuscript.

## Supplementary Figures

**Fig S1.**
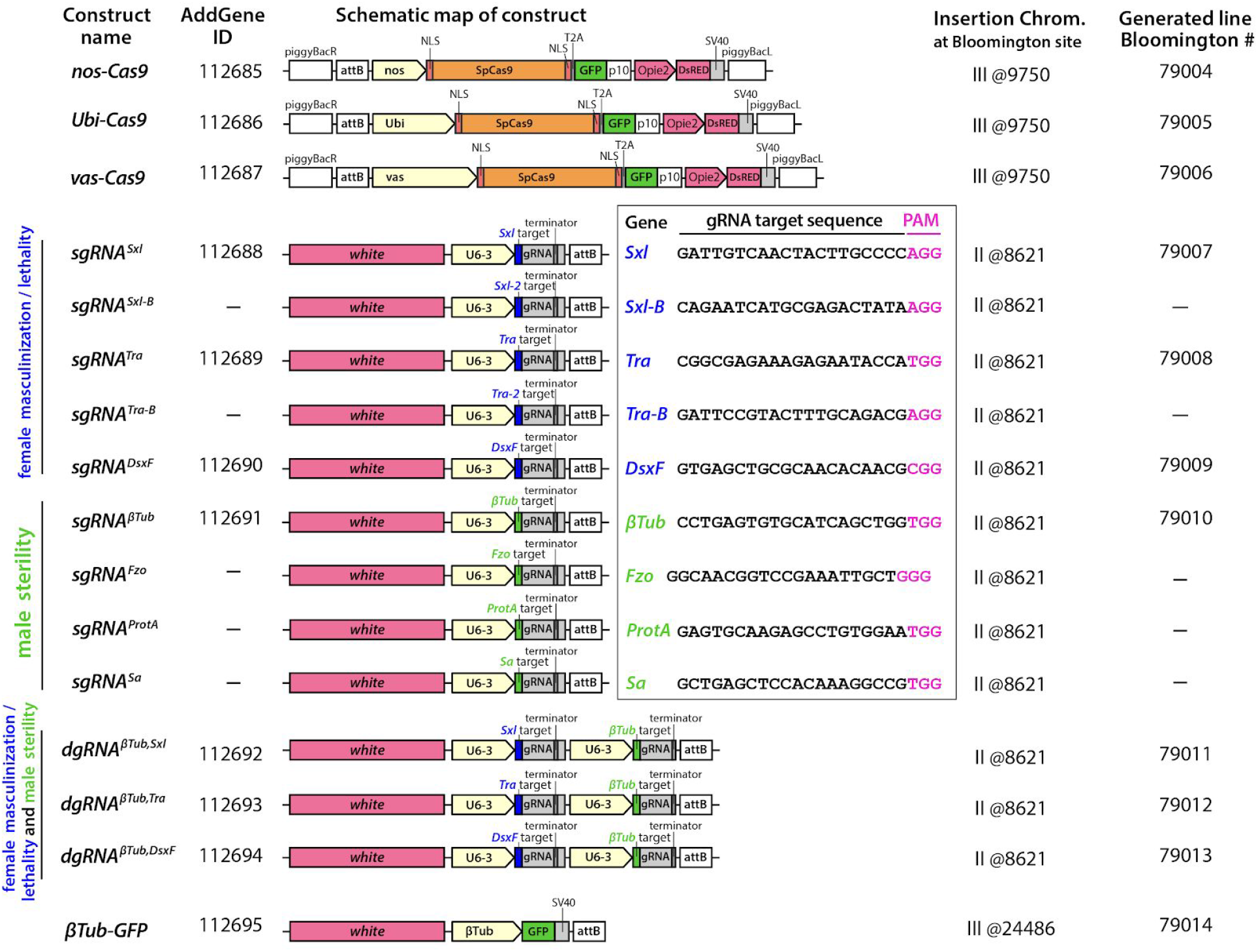
Genetic constructs and corresponding transgenic lines developed in the study. Schematics of all constructs engineered in this study, with functional constructs and flies deposited to Addgene.org and Bloomington Drosophila Stock Center, respectively. Gene names and gRNA target site sequences are presented in the box. The coding sequence of a *SpCas9* was flanked by two nuclear localization signals (NLS) at both ends and a self-cleaving T2A peptide with eGFP coding sequence at the C-end, serving as a visual indicator of Cas9 expression.

**Fig S2.**
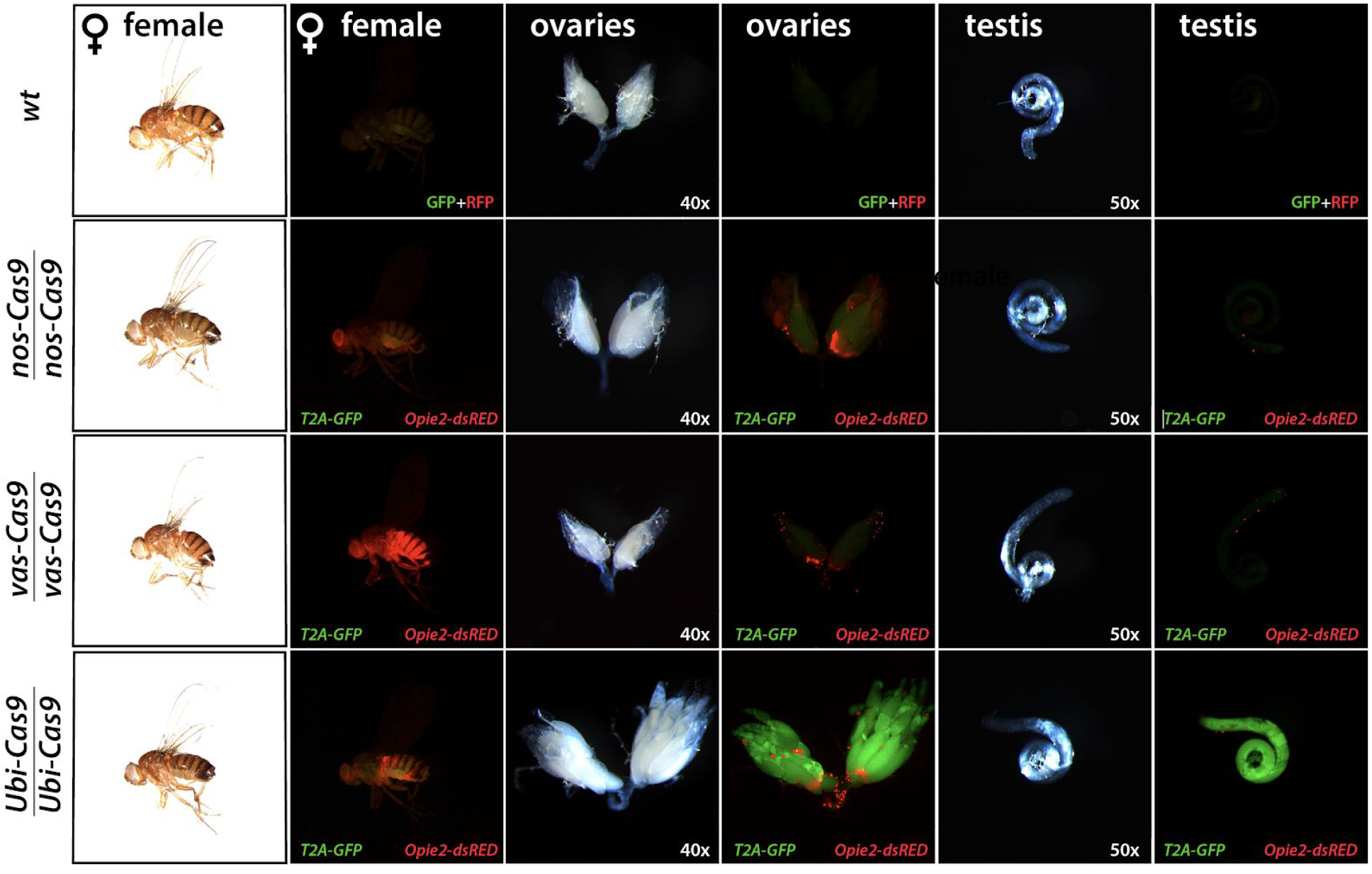
Characterization of three new homozygous lines expressing SpCas9. Three *Drosophila* lines supporting expression of *SpCas9* in strictly germline or germline together with somatic cells were developed. *Nanos-Cas9 (nos-Cas9), vasa-Cas9* (*vas-Cas9*), and *Ubiquitin-63E* (*Ubi-Cas9*) were inserted at the same site on the 3rd chromosome using φC31-mediated integration. *Opie2-dsRed* trasgene served as a transgenesis marker and a self-cleaving T2A-eGFP sequence, which was attached to the 3’-end of *SpCas9* coding sequence, provided an indicator of Cas9 expression (Fig. S1). Expression levels of dsRed and eGFP in each Cas9 line were compared to *wild type* (*wt*) flies. The Cas9-T2A-eGFP expression was mostly limited to female germline in *nos-Cas9* and *vas-Cas9* with a strong expression in *nos-Cas9. Ubi-Cas9* supported the strongest expression of Cas9, measured by eGFP, in both female and male germline, and in soma.

**Fig S3.**
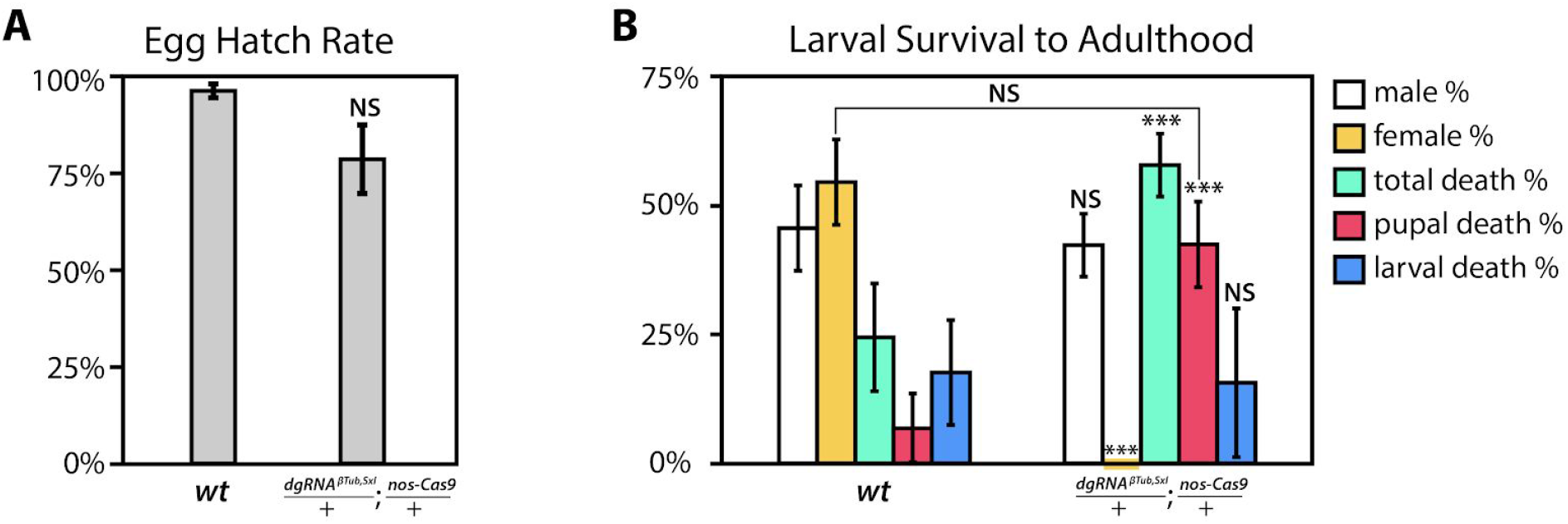
Determination of developmental stage of female lethality. (**A**) The percentage hatching rate estimated for *dgRNA^βTub,Sxl^/+; nos-Cas9/+* eggs generated by crossing homozygous *nos-Cas9* ♀ and *dgRNA^βTub-Sxl^ ♂* was not statistically different from that of the wild type (*wt*) eggs (Table S5). (**B**) Rates of different outcomes for hatched *dgRNA^βTub-Sxl^/+; nos-Cas9/+* larvae. Batches of 50 hatched larvae were raised to adults, their gender or developmental time of death was recorded (Table S6). The majority of additional larval deaths happened during a pupal transition, and the percentage of pupal death was not statistically different from the *wt* ♀ percentage. Statistical significance was calculated with a *t* tests assuming unequal variance. (*P <0.05^NS^, P* > 0.001***).

**Fig S4.**
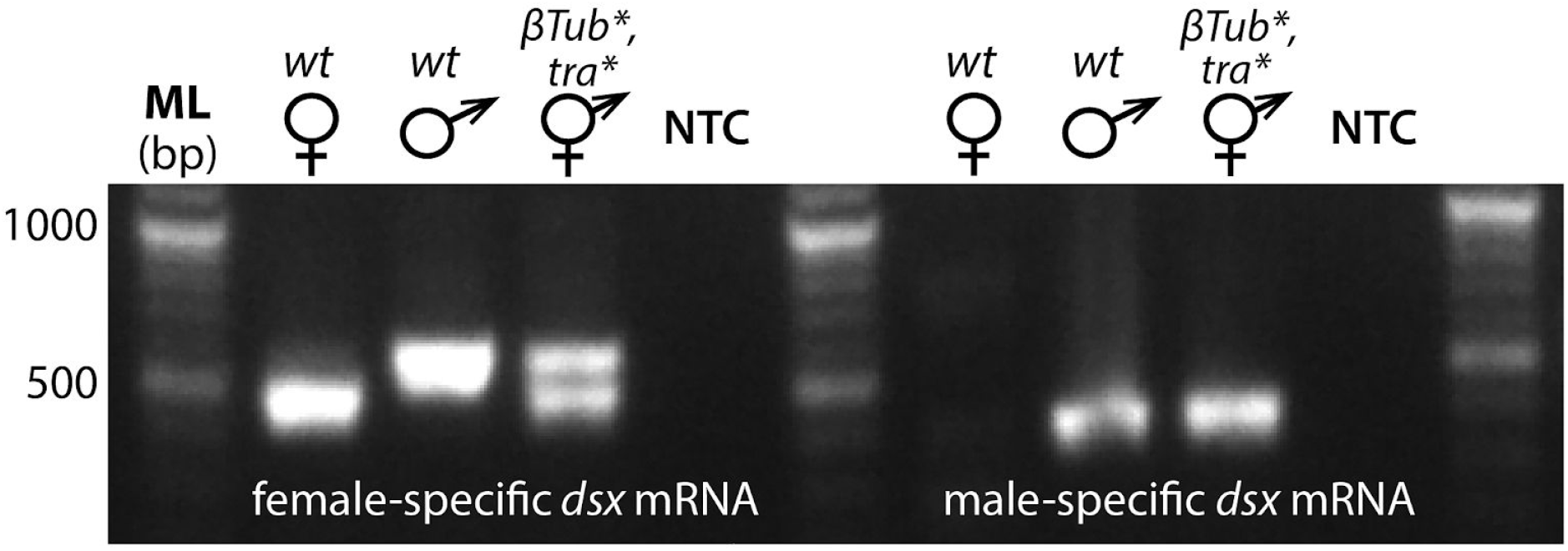
Both male and female splice variants of *Dsx* are expressed in *βTub, Tra* knockout intersexes. RT-PCR was used to assess female-specific and male-specific alternative splice variants of *dsx* comparing wild type (*wt*) females (♀), *wt* males (♂) and *dgRNA^βTub,Tra^/+; nos-Cas9/+* intersexes (*βTub*, Tra* ⚥*). Both female and male-specific *dsx* transcripts were identified in *βTub*, Tra*\* ⚥. Molecular ladder (ML) of double stranded DNA. No template control (NTC).

**Fig S5.**
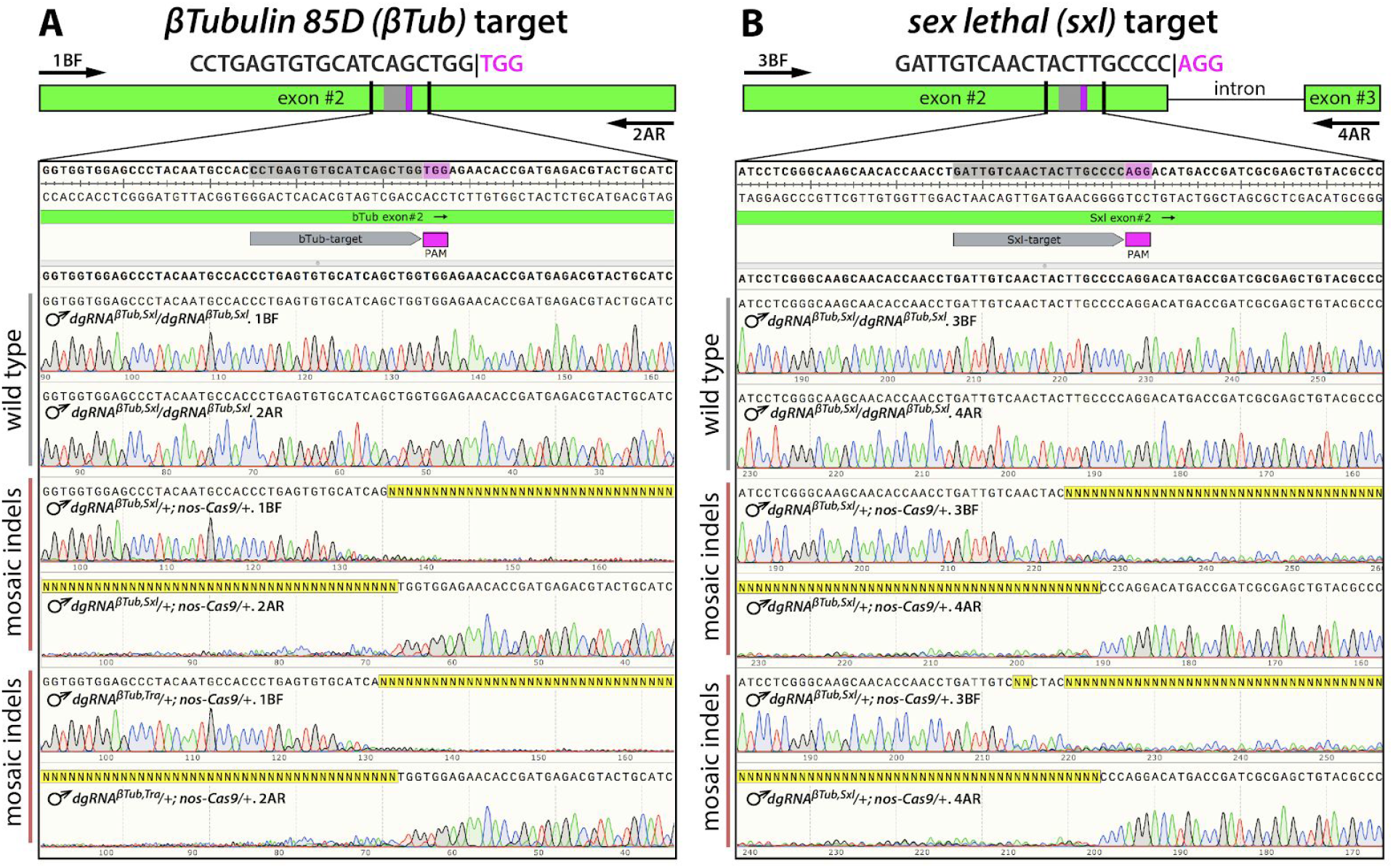
Biallelic mosaicism detected in the *βTub, Sxl* knockout flies. (**A**) Trans-heterozygous *dgRNA^βTub,Sxl^/+; nos-Cas9/+* sterile males (♂) had mosaic insertions / deletions (indels) precisely at the *βTub* target site. (**B**) Mosaic indels were also identified at the *Sxl* target site and likely caused the pupal lethality of trans-heterozygous females (Fig. S3). Diagrams on the top present positions of gRNA target sites and primers used for PCR relative to genetic structures of targeted genes. Sequence reads from both ends inferred diversity of templates that specifically localized at the sites targeted with gRNAs in the sterile ♂, while the wild type ♂ had single alleles without any sequence ambiguity.

**Fig S6.**
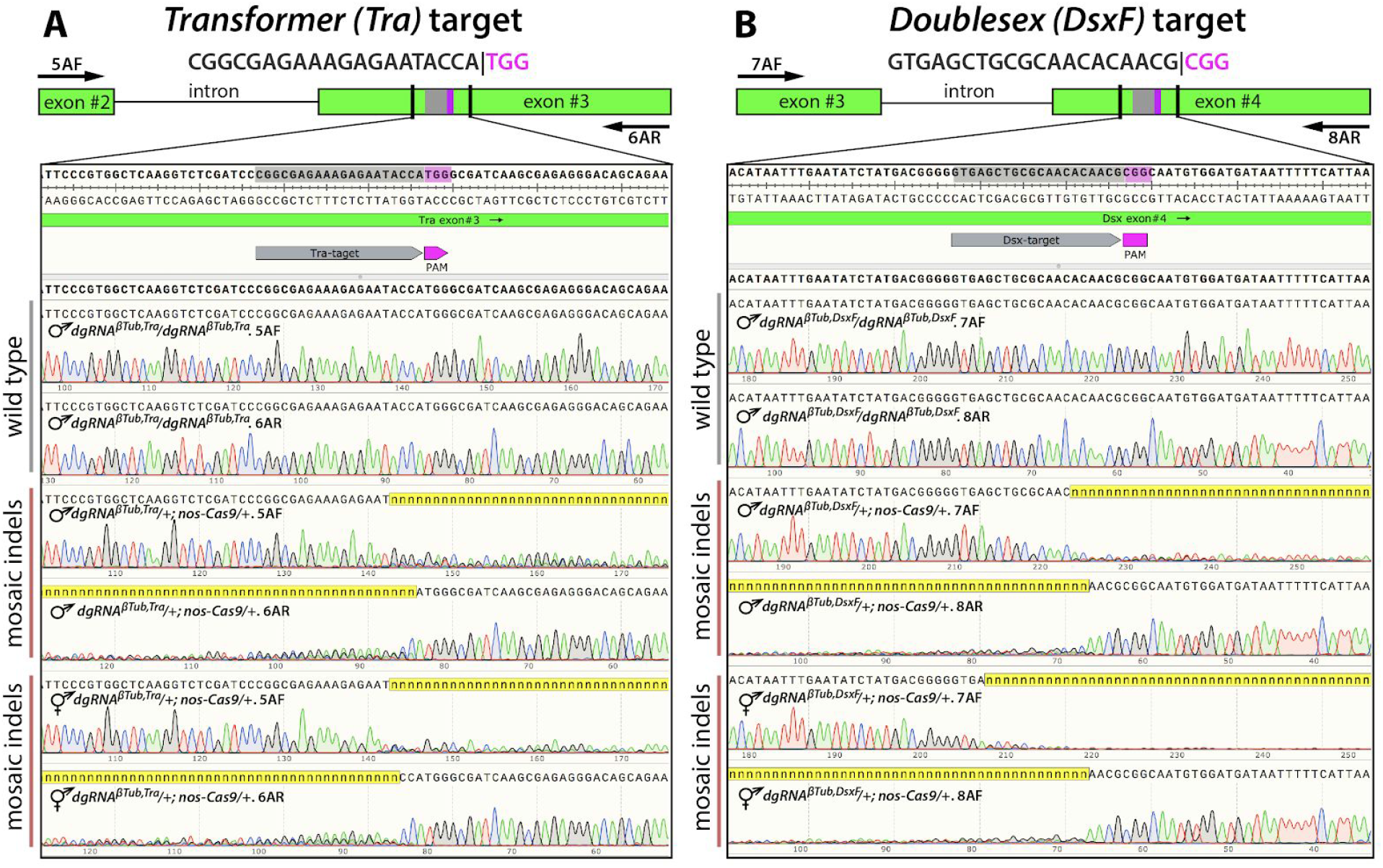
Biallelic mosaicism detected double knockout flies. (**A**) Trans-heterozygous *dgRNA^βTubJra^/+; nos-Cas9/+* sterile males (♂) and intersexes (⚥) had mosaic insertions / deletions (indels) located at the *Tra* site targeted by *dgRNA^βTub,Tra^* double guide RNAs (dgRNA). (**B**) Mosaic indels were identified at the *DsxF* site targeted by the *dgRNA^βTub,DsxF^* double gRNAs in *dgRNA^βTub,DsxF^/+; nos-Cas9/+* sterile ♂ and ⚥. Diagrams on the top show positions of gRNA targets and primers used for PCR relative to genetic structures of targeted genes. Sequence reads from both ends inferred diversity of templates that specifically localized at the sites targeted with gRNAs in sterile ♂ and ⚥, though the wild type ♂ had single alleles without any sequence ambiguity at both sites.

**Fig S7.**
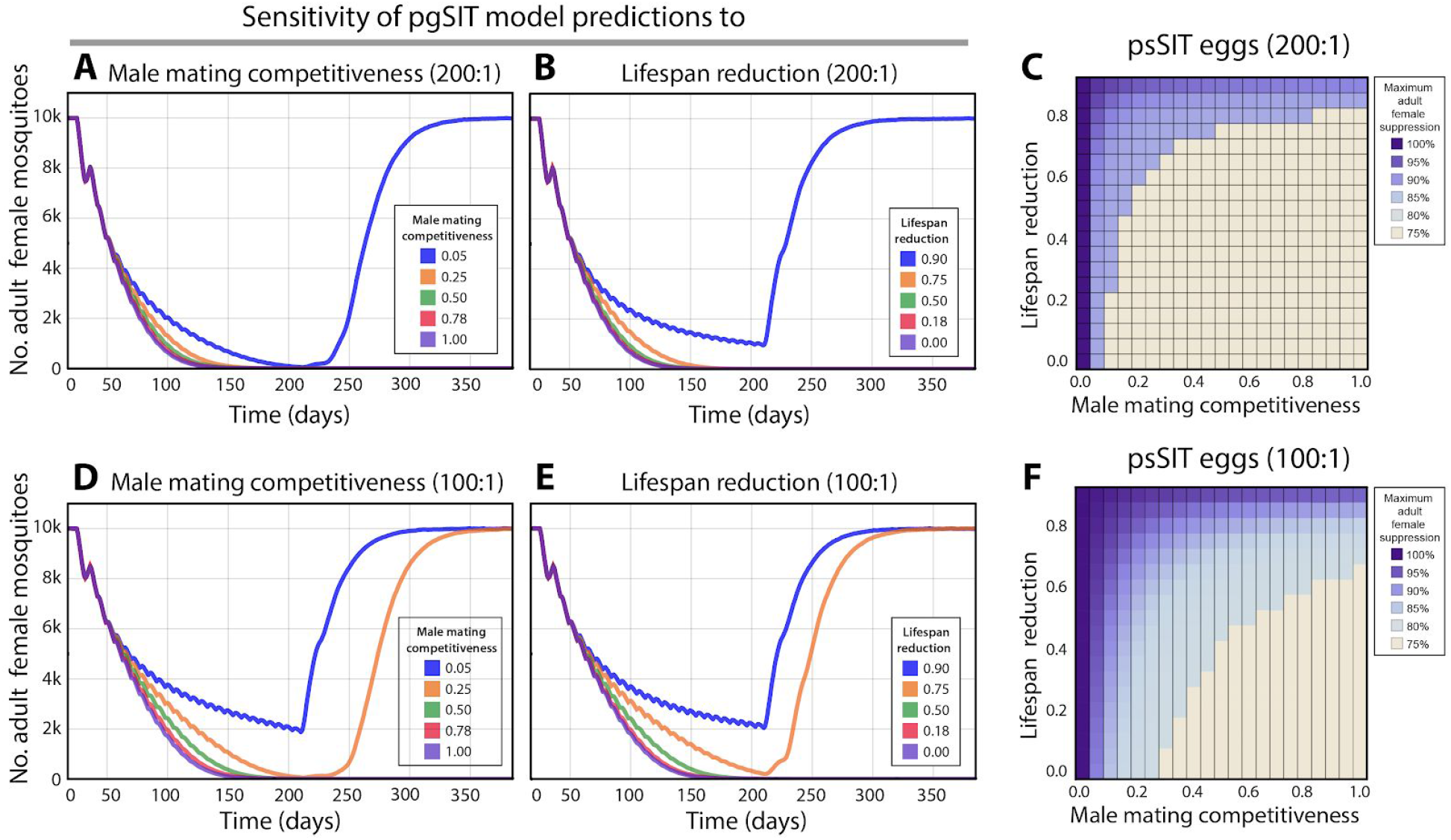
Sensitivity analysis for pgSIT system in *Aedes aegypti*. Sensitivity of pgSIT model predictions to male mating competitiveness, lifespan reduction and release ratio, keeping all other parameters constant as per Table S9. Model predictions were computed using 250 realizations of the stochastic implementation of the MGDrivE simulation framework ^61^ for a randomly-mixing *Ae. aegypti* population of 10,000 adult females. (**A**) For a weekly release ratio of 200 eggs per wild adult and keeping lifespan reduction due to the pgSIT construct constant at 18%, elimination can be reliably achieved for a male mating competitiveness of 25%; but not for 5%, as is the case for RIDL adult males. (**B**) For a weekly release ratio of 200 eggs per wild adult and keeping male mating competitiveness constant at 78%, elimination can be reliably achieved for lifespan reductions less than or equal to 75%. (**C**) Varying lifespan reduction and male mating competitiveness simultaneously suggests a wide range of parameter values for which local *Ae. aegypti* elimination could be reliably achieved (tan tiles) given a weekly release ratio of 200 eggs per wild adult. (**D**) For a weekly release ratio of 100 eggs per wild adult and keeping lifespan reduction due to the pgSIT construct constant at 18%, elimination can be reliably achieved for a male mating competitiveness of 50%; but not for 25%. (**E**) For a weekly release ratio of 100 eggs per wild adult and keeping male mating competitiveness constant at 78%, elimination can be reliably achieved for lifespan reductions less than or equal to 50%. (**F**) Varying lifespan reduction and male mating competitiveness simultaneously suggests a smaller yet still wide range of parameter values for which local *Ae. aegypti* elimination could be reliably achieved (tan tiles) given a weekly release ratio of 100 eggs per wild adult.

**Fig S8.**
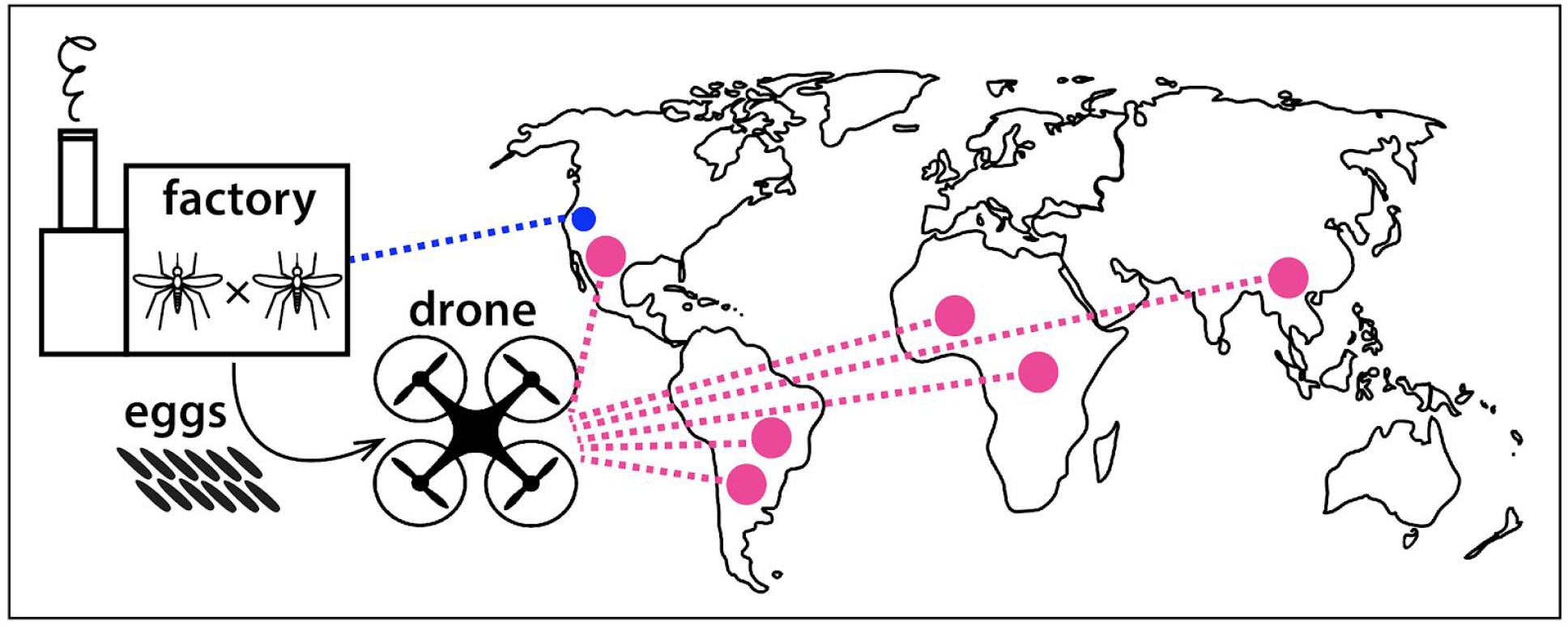
One factory (blue) producing pgSIT eggs for distribution and release (pink) at remote locations worldwide.

## Supplementary Tables

**Table S1:** Raw data from F_1_ progeny from the crosses between homozygous sgRNAs and homozygous Cas9. (Excel)

**Table S2:** Sequences of targeted sites: indels are found in transheterozygous flies. (Excel)

**Table S3:** Raw data from F_1_ progeny from the crosses between homozygous dgRNAs and homozygous Cas9. (Excel)

**Table S4:** Characterization of masculinized females (intersexes). (Word)

**Table S5:** Hatching rate of *gRNA^bTub-Sxl^/+; nos-Cas9/+* embryos. (Excel)

**Table S6:** Female pupal lethality in *dgRNA^bTub,Sxl^/+; nos-Cas9/+* embryos. (Excel)

**Table** S7: Raw data from F_1_ progeny from the crosses between homozygous dgRNAs and heterozygous Cas9. (Excel)

**Table S8:** Competitiveness of *dgRNA^bTub,Sxl^/+; nos-Cas9/+* males. (Excel)

**Table S9:** Parameters used in *Aedes aegypti* population suppression model. (Word)

**Table S10:** Primers used in this study. (Word)

